# Spatiotemporal discrimination in attractor networks with short-term synaptic plasticity

**DOI:** 10.1101/296905

**Authors:** Benjamin Shlaer, Benjamin Ballintyn, Paul Miller

## Abstract

We demonstrate the ability of a randomly connected attractor network with dynamic synapses to discriminate between similar sequences containing multiple stimuli and suggest such networks provide a general basis for neural computations in the brain. The network is based on units representing assemblies of pools of neurons, with preferentially strong recurrent excitatory connections within each unit. Such excitatory feedback to a unit can generate bistability, though in many networks only under conditions of net excitatory input from other units. Weak interactions between units leads to a multiplicity of attractor states, within which information can persist beyond stimulus offset. When a new stimulus arrives, the prior state of the network impacts the encoding of the incoming information, with short-term synaptic depression ensuring an itinerancy between sets of active units. We assess the ability of such a network to encode the identity of sequences of stimuli, so as to provide a template for sequence recall, or decisions based on accumulation of evidence. Across a range of parameters, such networks produce the primacy (better final encoding of the earliest stimuli) and recency (better final encoding of the latest stimuli) observed in human recall data and can retain the information needed to make a binary choice based on total number of presentations of a specific stimulus. Similarities and differences in the final states of the network produced by different sequences lead to predictions of specific errors that could arise when an animal or human subject generalizes from training data, when the training data comprises a subset of the entire stimulus repertoire. We suggest that such networks can provide the robust general purpose computational engines needed for us to solve many cognitive tasks.

## Introduction

Life is a sequence of interconnected events, such that our optimal response to one event is highly impacted by preceding events. For example, our actions following the sound of a fire alarm should be very different if we had just received a message about a forthcoming alarm test, from our actions following the same sound if we had just seen smoke emanate from a nearby chemistry laboratory. Moreover, the impact of auditory stimuli, most clearly noticeable in music or speech, is significantly affected by the stimuli preceding it. A musical note rarely sounds “pleasant” or “unpleasant” alone, yet can do so when expectations are set by an ongoing melody. Our response to someone yelling the word “run” is very different if the word “run” is preceded by the two words “do not”. Therefore, a key aspect of cognition is to enable our response to any stimulus to depend upon the preceding pattern of inputs. Fundamental to such an ability is the need for the pattern of neural activity in the underlying circuits to depend not just on the current stimulus, but on the entire sequence. In this paper, we propose one general method, based on point-attractor states, that the brain might use for achieving such sequence-dependent activity.

### Circuits with multiple point-attractor states

Networks of recurrently connected units are capable of producing a diversity of distinct point-attractor states [1]. The component units of these networks can be single neurons, or groups of correlated, similarly responsive neurons. The stable activity states they produce are called ‘attractors’, because when a stimulus causes the network’s activity to become similar to, or ‘nearby’ the attractor, the internal dynamics of the network cause that activity to shift toward (or be ‘attracted to’) the attractor state. The states are point attractors, because they are each described by a single, distinct set of firing rates of all the units. The set of rates can be represented as a single point in the high-dimensional space where each axis corresponds to a single unit’s rate. Such attractor states can arise when the neurons in a network change their connections via Hebbian synaptic plasticity [2, 3]. In autoassociative networks, such as the Hopfield network [4], such plasticity allows for the learning and long-term storage of input patterns, which can be reconstructed later from partial or corrupted versions of the prior stimuli. In this scheme, after learning, attractor states of the network represent learned representations of the presented stimuli.

Modeling studies have also shown that networks with point-attractor states can underlie short-term memory and decision-making [5–7]. In vivo experiments have provided support for these ideas [8–14]. In recurrent networks with strong self-excitatory connections, units are capable of maintaining their firing state long after the presentation of a stimulus. Such behavior is known as bistability since the neuron has two stable firing rates--quiescent and active-- in the absence of stimulus. In such a network, the activity or lack thereof within a particular group of cells forms a basis for short-term memory of prior inputs [15].

Recurrent networks with random excitatory connections have been shown to have particularly useful properties [16]. Random network connections in a recurrent network endow the neurons with ‘mixed-selectivity’ for combinations of stimuli, a property of neurons observed in vivo [17, 18] and important for solving linearly non-separable tasks (such as the Exclusive-Or) [3].

If the inputs to neurons in an attractor network are relatively strong compared to the recurrent feedback, then when the network receives a new input its activity will switch to the new attractor state corresponding to the new input. In such an event, information about the prior input is lost. Alternatively, if the input is relatively weak then it can be insufficient to drive the activity away from the prior attractor state, and only information about the initial stimulus is retained. We suggest that this parameter dependence, which leads to either recency (strongest memory for the last stimulus) or primacy (strongest memory for the first stimulus) might be the basis of these observed recall phenomena [19]. Given individual human subjects exhibit both primacy and recency in a single task [20], we investigate whether both effects can arise in a single heterogeneous network.

Evidence for quasistable point attractors in neural circuits in vivo arises from the observation of rapid transitions between relatively stationary states of coordinated network activity. Such rapid transitions can occur in the absence of a stimulus, or when the presented stimulus is constant or gradually ramping. For example, Mattia et al. [21] found that in a visuomotor decision-making task, multi-unit recordings in pre-motor cortex revealed that groups of neurons transition suddenly between distinct activity states, forming a series of network states which ultimately settle into a stereotyped network activity pattern that predicts future movement. Additional in vivo evidence comes from rat and mouse gustatory and auditory cortices [7, 11, 22, 23]. Both sensory cortices show series of abrupt transitions between discrete network states, whose timing varies from trial-to-trial, such that the transitions may be obscured by typical analyses with cross-trial averaging.

Evidence of point-attractors has also been observed to govern place field remapping in the hippocampus [24]. Place cell populations responded to gradual changes in the environment with a rapid transition in network state, thought to represent environmental context. Taken together, these studies provide extensive evidence for point-attractor dynamics in vivo [14].

### Sequence-dependent memory

In order for a network to possess sequence-dependent memory, it must exhibit both of the following characteristics. First, it must respond to an external stimulus in a manner that discriminates between all possible incoming stimuli. Second, it must do so while retaining information about the pre-stimulus state of the network. An added complication is that this behavior must succeed in the presence of realistic noise fluctuations.

As demonstrated previously [25], the dynamics of synaptic depression can enable a randomly connected attractor network to respond to new stimuli while retaining information about the pre-stimulus state. Once a stimulus is removed, the stable state of the network (a fixed point) can depend both on the most recent stimulus and on the state of the network prior to that stimulus. Repeating the process in this manner with successive stimuli can lead to a network state that depends on the properties of an entire sequence of stimuli. We showed that these networks can encode in distinct states the amplitude, duration, and number of stimuli presented to the network in a temporal sequence. The distinct activity states made use of the high-dimensional space of neural firing rates [26], so that the different stimulus properties are not combined together as they would be in standard models, such as neural integrators [27–31], which have low dimensionality [32]. In the successful models of ours, each stimulus was encoded as an input of identical magnitude to all cells, with different stimuli distinguished by the duration and amplitude of that input. Here we extend this result to show that versions of the same network can encode temporal sequences of different stimuli, in which the stimuli are distinguished according to which cells they activate.

We perform two discrimination tasks with the network. In the first, we consider sequences comprised of different patterns of two distinct stimuli, separated by short pauses, with six stimuli combined into the sequence. This task is motivated by an experiment [33] in which a mouse is trained to turn left or right at the junction of a T-shaped maze according to whether it sees more visual cues on its left or right side when approaching the junction. Such a task can be solved by a circuit which takes the difference between the two counters or integrators (one for the left, one for the right) requiring only a one-dimensional representation of stimulus history (a single scalar). However, the authors found that neural activity stepped through sequences of states that depended on the entire preceding sequence of stimuli. The observed dynamics is more suggestive of the distributed coding present in a high-dimensional representation of stimulus history. Such a high-dimensional representation arises in the randomly connected network of bistable units that we consider here.

The second task requires discrimination between sequences of seven distinct stimuli presented without repetition. We test the network’s ability to discriminate between different permutations of the seven stimuli, and to what extent the final state of the network could be said to encode each one of the presented stimuli, compared to stimuli that had not been presented. The work is motivated by experiments on free recall of lists of presented words. In those experiments, better recall of the first one or two words in the list (primacy) and/or better recall of the final one or two words in the list (recency) is typically found [19, 34–36].

We will test the extent to which our network possesses primacy and/or recency by comparing how much the final state is impacted by changes in the first stimulus or the last stimulus, compared to changes in the middle of the sequence of presented stimuli. A network with strong self-excitation and strong cross-inhibition may, after encoding one stimulus, suppress any change in its activity when later stimuli are presented. Such a network would produce strong primacy, with its final state being very similar to its state following just the first stimulus. In this case, its final state would be more strongly affected by a swap of the first two stimuli than by a swap of later stimuli. Conversely, a network dominated by the external input, rather than the internal feedback, is more likely to shift its activity entirely to match the latest input pattern. Such a network would exhibit recency, with its final activity most similar to the pattern produced by the latest input alone. In this case, the network’s final state would be more altered by a swap of the last two items in the sequence than by a swap among any of the prior items in the sequence. Strong noise currents contribute only to recency, since they can eventually wash out information stored earlier in the network.

The transient weakening of self-interactions, due to synaptic depression, endows the network with a propensity for self-avoiding trajectories in the space of firing rates, a useful feature for discriminating temporal information about sequences of stimuli [37]. Since units that were recently on (*τ*_*D*_ = 500*ms*) are less likely to become active while the network is evolving towards a fixed point, the network preferentially takes large steps in activity space following new stimuli. Large steps mean a large number of new fixed points that are within range. This facilitates discrimination of temporal sequences of stimuli because confusion occurs whenever two distinct initial states can be brought to the same final state via presentation of identical stimuli.

## Methods

We summarize the model and stimuli in Table 1 below.

**Table 1.**
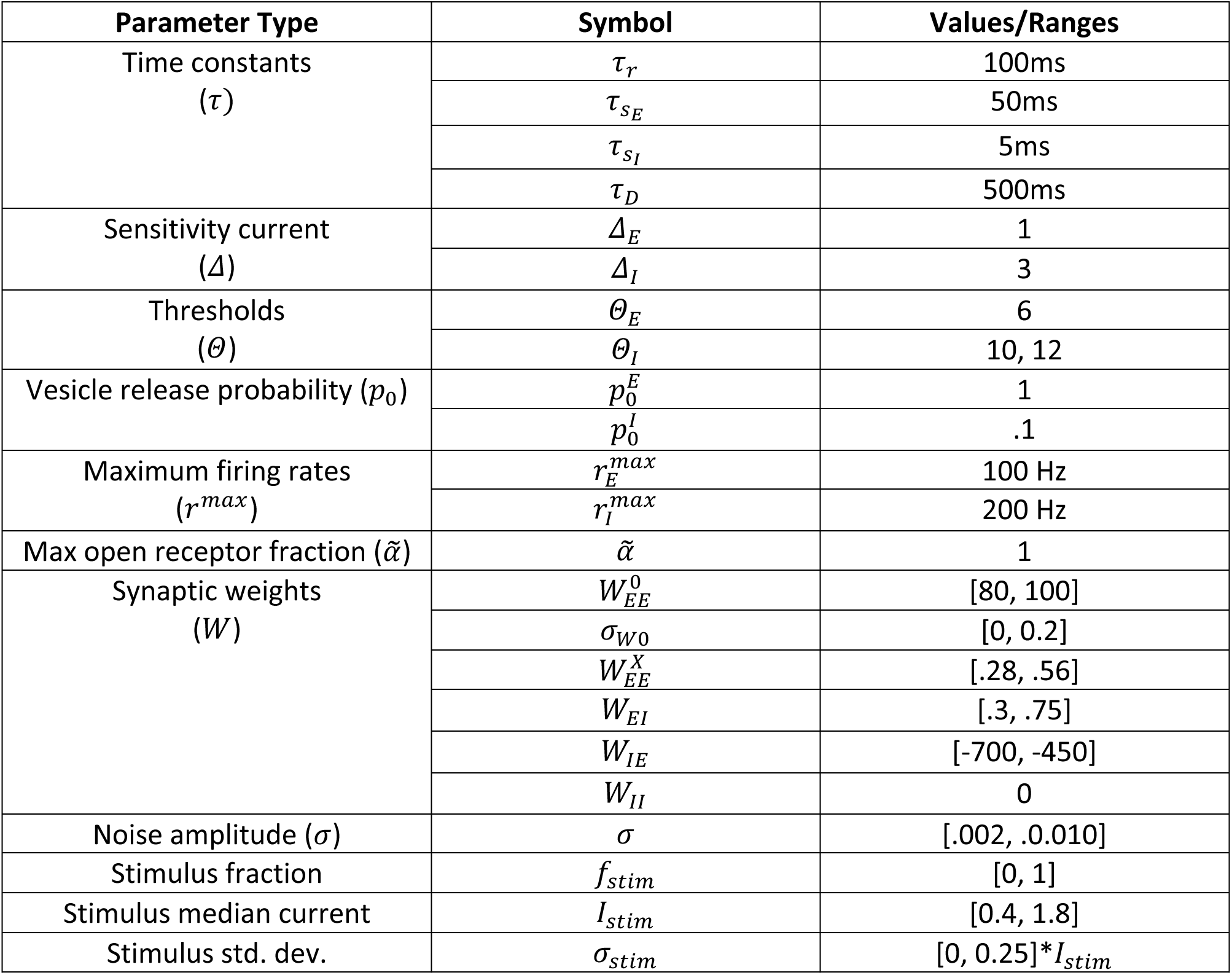
Model parameter ranges.

| Parameter Type | Symbol | Values/Ranges |
| --- | --- | --- |
| Time constants<br>($\tau$) | $\tau_r$ | 100ms |
| | $\tau_{sE}$ | 50ms |
| | $\tau_{sI}$ | 5ms |
| | $\tau_D$ | 500ms |
| Sensitivity current<br>($\Delta$) | $\Delta_E$ | 1 |
| | $\Delta_I$ | 3 |
| Thresholds<br>($\theta$) | $\theta_E$ | 6 |
| | $\theta_I$ | 10, 12 |
| Vesicle release probability ($p_0$) | $p_0^E$ | 1 |
| | $p_0^I$ | .1 |
| Maximum firing rates<br>($r^{max}$) | $r_E^{max}$ | 100 Hz |
| | $r_I^{max}$ | 200 Hz |
| Max open receptor fraction ($\tilde{\alpha}$) | $\tilde{\alpha}$ | 1 |
| Synaptic weights<br>($W$) | $W_{EE}^0$ | [80, 100] |
| | $\sigma_{W0}$ | [0, 0.2] |
| | $W_{EE}^X$ | [.28, .56] |
| | $W_{EI}$ | [.3, .75] |
| | $W_{IE}$ | [-700, -450] |
| | $W_{II}$ | 0 |
| Noise amplitude ($\sigma$) | $\sigma$ | [.002, .010] |
| Stimulus fraction | $f_{stim}$ | [0, 1] |
| Stimulus median current | $I_{stim}$ | [0.4, 1.8] |
| Stimulus std. dev. | $\sigma_{stim}$ | [0, 0.25]*$I_{stim}$ |

Summary of methods.

### A. Model Summary

|  |  |
| --- | --- |
| Populations | $N_E = 100$ excitatory, $N_I = 1$ inhibitory |
| Connectivity | E-to-E: all-to-all with random strength, strong self-excitation |
| Neuron model | Firing rate model |
| Synapse model | Single exponential with synaptic depression |
| Plasticity | No long-term plasticity |
| Input | Sequences of spatially variable fixed duration square pulses |
| Measurements | Long lived (meta stable) firing rates following stimulus sequences |

### B. Populations

| Name | Type | Number |
| --- | --- | --- |
| Excitatory (E) | Firing rate model | $N_E = 100$ |
| Inhibitory (I) | Firing rate model | $N_I = 1$ |

### C. Connectivity

| Name | Source | Target | Distribution |
| --- | --- | --- | --- |
| $W_{EE}^0$ | E | Self | Uniform iid $\in [(1 - \sigma_0), 1]$ $W_{EE}^0$ |
| $W_{EE}^X$ | E | Other E | Uniform iid $\in [0, W_{EE}^X]$ |
| $W_{EI}$ | E | I | Fixed at $W_{EI} > 0$ |
| $W_{IE}$ | I | E | Fixed at $W_{IE} < 0$ |
| $W_{II}$ | I | I | Fixed at 0 |

### D. Neuron and synapse model

|  |  |
| --- | --- |
| Name | Firing rate model with synaptic depression |
| Type | Leaky integrate-and-fire with exponential conductance input |
| Input current | $I_j(t) = \sum_i s_i(t) W_{ij} + I_j^{app}(t) + \sigma \eta_j(t)$ |
| Firing rate | $\tau_r \frac{dr_j(t)}{dt} = -r_j(t) + \frac{r^{max}}{1 + e^{(\theta_j - I_j(t))/\Delta_j}}$ |
| Synaptic gating variable | $\tau_s \frac{ds_i(t)}{dt} = -s_i(t) + \tilde{\alpha} p_0 r_i(t) D_i(t) \tau_s (1 - s_i(t))$ |
| Depression Variable | $\tau_D \frac{dD_i(t)}{dt} = 1 - D_i(t) - p_0 r_i(t) \tau_D D_i(t)$ |

### E. Plasticity

There is no long-term plasticity in this model, only short-term synaptic depression.

### F. Stimuli

Stimuli are spatially variable synchronized square pulses of median current (*I*_stim_) to a fraction of excitatory units (*f*_stim_). Each unit within this fraction receives current with amplitude independently drawn from a log-normal distribution with standard deviation (σ_stim_). A summary of the stimulus sequences used for each task is given in the table below.

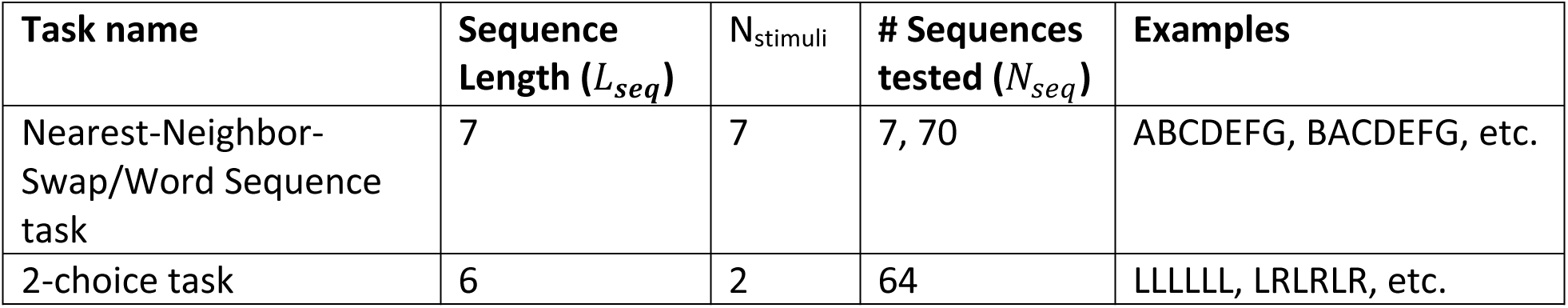

### G. Measurements

The primary measurement from which our analysis is derived is the high-dimensional state of the network following a given stimulus sequence. We measure the network state by taking the time-averaged firing rates of excitatory units following the final stimulus in a sequence.

#### 1) Firing rate model with synaptic depression

The basic unit of our network represents a pool of tightly coupled neurons. Their strong mutual interaction allows us to model this pool as a single unit, which can exhibit bistability with sufficient input current. The *i*^*th*^ unit is characterized by a single firing rate *r*_*i*_(*t*), dependent upon its net input current via a sigmoidal *f*-I curve,

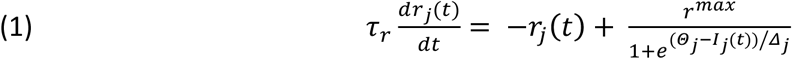

where *τ*_*r*_ = 10*ms* is the firing rate time constant, *Θ*_*i*_ is the threshold current, and *Δ*_*i*_ is the sensitivity current (controlling the slope of the *f*-I curve). The input current to the *i*^*th*^ cell-group is given by,

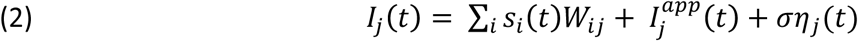

where *ση*_*j*_(*t*) is a white noise current with standard deviation *σ*, 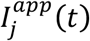 is the external applied current (stimulus), *s*_*i*_(*t*) is the dimensionless synaptic gating variable from the *i*^*th*^ group of cells, and *W*_*ij*_ is the connectivity matrix. The synaptic gating variable obeys

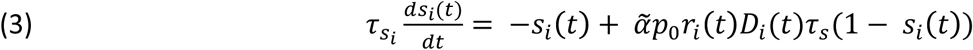

where the synaptic time constant *τ*_*s*_ describes the decay time of *s*_*i*_ following a significant decrease in firing rate. Excitatory units had a *τ*_*s*_*E*__ = 50*ms* while the single inhibitory unit had a *τ*_*s*_I__ = 5*ms*. The increase in *s*_*i*_(*t*) toward unity is proportional to the firing rate *r*_*i*_(*t*) and the dimensionless depression variable *D*_*i*_(*t*). The fraction of open receptors in response to maximal vesicle release is *ᾶ* which we set to unity. The depression variable obeys,

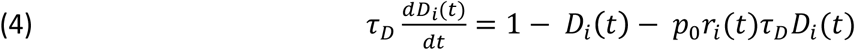

where *τ*_*D*_ = 500*ms.*

The ranges of parameters used in our simulations are summarized in the table below.

#### 2) Stimuli

Each stimulus consists of a square current pulse of duration 250ms to a randomly chosen fraction of excitatory units. We will describe these stimuli as spatial patterns, each consisting of *N*_*E*_ * *f*_*stim*_ currents independently drawn from a log-normal distribution. Stimulus sequences consist of *L*_*seq*_ of these 250ms current pulses delivered in some order separated by 1.5 seconds. This leaves a gap of 1250ms during which the activity of the network may evolve without external stimulus toward a new fixed-point. The single inhibitory unit receives no external current.

#### 3) Measurements

After a complete sequence of stimuli is presented, we wait 1150ms before beginning to average the firing rates. The average is taken over 100ms, i.e., between 1150ms and 1250ms after the final stimulus offset. This average firing rate is then used in all decoding efforts.

#### 4) Trials

For each of the two discrimination tasks, we perform the following steps:

1. Select network parameters (5 connectivity matrix parameters, stimulus amplitude, stimulus fraction, stimulus variance, and noise level).

2. Generate a random instance of the network using these parameters

3. Generate the random input currents representing each stimulus sequence

4. Present the stimulus sequences to the network and record the final network state after each sequence.

5. Evaluate the network’s ability to discriminate between sequences and its ability to recall individual stimuli.

The random seed used for network and stimulus generation is distinct for each parameter combination, as is the random seed used to generate the noise within the units.

#### 5) Decoding discrimination

The ability of a network to discriminate between stimuli depends on how often two distinct stimuli lead to the same network activity. Due to the presence of noise, which broadens the distributions associated to each stimulus, estimating this quantity is numerically intractable for all but the smallest networks. In cases where the number of distinct stimulus sequences is small, cluster analysis can be used. A more generally applicable method, which works both in large networks and for those receiving many stimuli, is to train a decoder with the mean “target” response to each stimulus sequence. Then any test response can be compared with the previously measured target responses to see which is closest.

Specifically, to quantify a network’s ability to discriminate between sequences, we first prepared the network in a quiescent initial state. We then allowed the network to evolve with no external stimuli for at least five seconds, and used the resulting state as our initial condition for all subsequent stimulus presentations. We presented each stimulus sequence ten times to establish the supervised mean (“target”) response. We then presented each stimulus sequence ten additional times, and assigned to each a prediction based on the minimum L1-norm (shortest taxicab-distance) in firing rate space between this instance and the target response. Once we have determined the predicted sequence for each of the ten repetitions of each of the *N*_*seq*_ sequences, we can construct a confusion matrix (Figure 2), where element (*i,j*) represents the probability that actual presentation of a test sequence, *j*, results in the network predicting that the target sequence, *i*, was presented. Hence each column of the confusion matrix sums to unity, and all of the information is contained in the off-diagonal elements, known as the error-rates.

**Figure 1.**
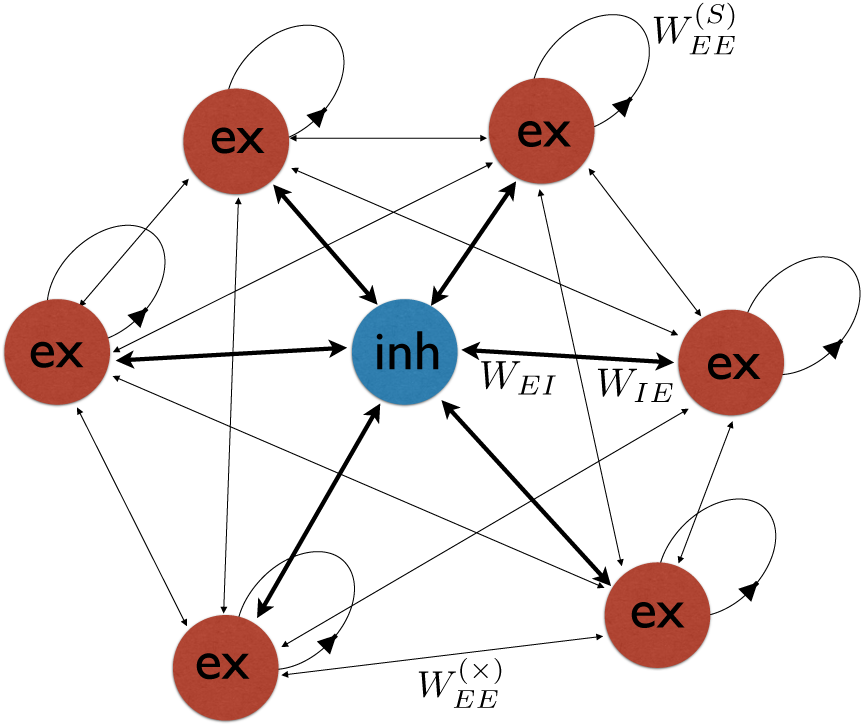
Schematic view of network. Excitatory self-connections render each excitatory unit bi-stable.

**Figure 2.**
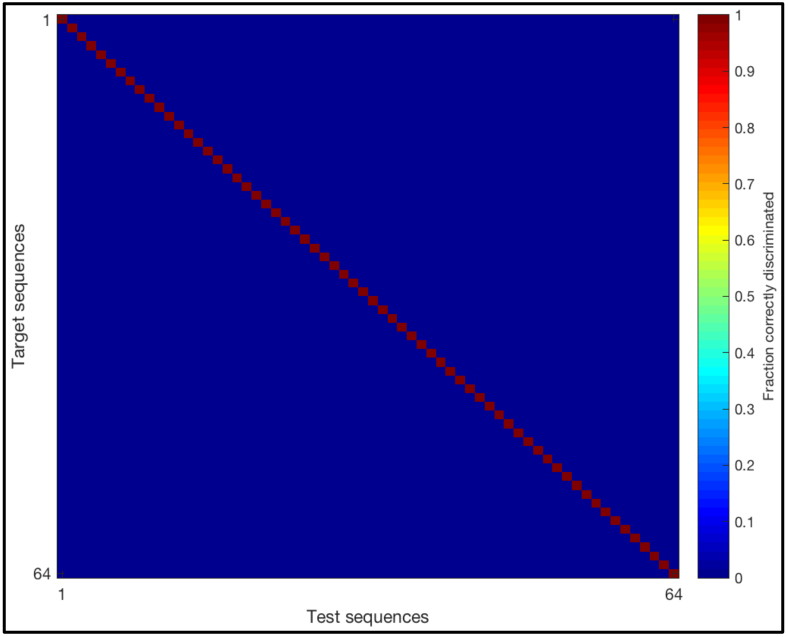
Confusion matrix for a particular network. Entry **(*i,j*)** gives the fraction of times test sequence j was identified as target sequence i. This particular network 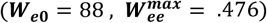 achieved a perfect discrimination accuracy **(*κ* = 1)**.

#### 6) Measuring performance

##### 6.1 Sequence Discrimination ability

We use a confusion matrix ***C*** to quantify the discrimination performance of the network. This matrix can be converted into several interesting scalar quantities. The total discrimination ability (*κ*) is defined as the fraction of times the network correctly classified the mean activity from a sequence, linearly rescaled such that a score of zero is obtained for discriminating no better than chance, and a score of unity is a perfect score:

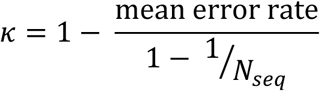

where *N*_*seq*_ is the number of distinct sequences. Although it is possible for a network to perform worse than chance, we clip all *κ* values to be ∈ [0,1].

##### 6.2 Primacy and recency

We used 2 methods to quantify the effects of primacy and recency in each network. In the first, we train perceptrons to indicate, based on the final network state, whether stimulus type X was present in serial position *sp* of each sequence. In the second method, we extract the network’s ability to discriminate changes in stimuli at a particular position, using a metric (*κ*_*sp*_) derived from the confusion matrix.

In the first method, we first collect the final network states of all target-trials (10 target trials for each sequence) into a *N*_*E*_×(*N*_*trials*_ * *N*_*seq*_) matrix ***X***_***train***_, where the final state of each trial is a column in the matrix. The next step depends on which model task we are analyzing. For the 2-choice task, since each sequence consists of 6 stimuli of only 2 types, for each network we train 6 perceptrons (*P*_1_,*P*_2_,…,*P*_6_) on ***X***_***train***_ to indicate, based on the final network state, whether the stimulus in serial position *SP* was of type 1 or 2. We then evaluate each perceptron *P*_*sp*_ on a separate test dataset, *X*_*test*_, which also contains 10 trials for each sequence, and obtain a “recall” accuracy, *a*_*sp*_, for each serial position. The process for the word sequence task is similar, however because there are 7 different stimulus types and 7 stimuli in each sequence, we train 7 perceptrons for each serial position (1 perceptron per stimulus type per serial position) for a total of 49 perceptrons where each is trained on ***X***_*train*_ to indicate whether stimulus type *i* was present in serial position *sp* or not. Then, for each serial position, we can compute 7 accuracies (1 for each stimulus type) and we set *a*_*sp*_ to be the mean of these 7 accuracies.

In the second method, we extract from the confusion matrix the sub-abilities for the network to discriminate based on the serial positions of the sequence differences (for example, ABCDEFG and ABDCEFG differ in their 3rd and 4th serial positions). For sequences which differ in (at least) their *sp*^*th*^ stimulus, where 1 ≤ *sp* ≤ *L*_*seq*_ and where *L*_*seq*_ is the number of stimuli in a complete sequence, we can measure

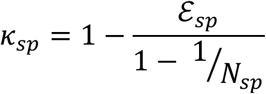

where *ε*_*sp*_ (*sp* -error rate) is the mean rate at which a sequence is misclassified as one which differs in its *sp*^*th*^ serial position and *N*_*sp*_ is the number of sequences which differ in their *sp*^*th*^ serial position. Note that *ε*_*sp*_ is straightforward to extract from the confusion matrix and a simple method for rescaling any performance metric of the confusion matrix *F*(*C*) is to use

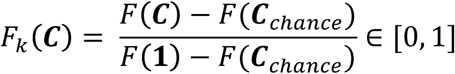

where the identity matrix **1** is the error-less confusion matrix, and ***C***_*chance*_ is the uniform matrix with all entries equal to 1/*N*_*seq*_, representing the confusion matrix obtained by guessing uniformly.

These “recall” abilities (*a*_*sp*_ in the first method and ***κ***_*sp*_ in the second) for each serial position can then be plotted (on an evenly spaced interval from x = 0 for the first stimulus to x = 1 for the last stimulus) and fitted with a quadratic (of the form: *ax*^2^ + *bx* + *c*). A primacy effect would show up on this plot as a higher “recall” ability for the earliest stimuli (with a negative sloping quadratic at x = 0) while recency would show up as a higher “recall” ability for the latest stimuli (with a positive slope at x = 1). We therefore define the ***primacy score*** of each network to be the negative of the derivative of the quadratic at x = 0:

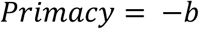

and the ***recency score*** to be the derivative at x = 1:

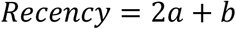

We set the primacy score to be the negative of the derivative so that larger positive numbers of each score indicate more of each respective phenomenon.

#### 7) Left/Right sequence discrimination and evidence accumulation

The first of the two discrimination tasks is based on the experiment of Morcos and Harvey [33]. The network is presented with a sequence of six cues of 2 types (modeling visual stimuli in the left or right visual field) and must make a binary decision based on the number of each type of stimuli (choose left if there are more left stimuli and right if there are more right stimuli). Hence, *N*_*types*_ = 2, *N*_*stim*_ = 6, *N*_*seq*_ = 2^6^ = 64. The network must keep track of how many of each stimulus type there is in the sequence and make a decision (turn left or right). In order to model the binary decision in the task we trained a perceptron to make the correct binary decision given a stimulus sequence (e.g. if a sequence contained more of stimulus type 1 than 2 the perceptron should output 0, otherwise if there were more of stimulus type 2 it should output 1). The perceptron was trained on a matrix of final network states (similar to ***X***_***train***_ but with the final states of sequences containing an equal number of both stimuli omitted). The perceptron was then evaluated on a test dataset *X*_*test*_ and its accuracy (*a*_*decision*_ = fraction of correct choices) calculated.

#### 8) Seven-word list discrimination

The second of the two discrimination tasks models word sequence recall in humans. The task consists of presenting sequences of 7 distinct stimuli and testing whether the sequences can be discriminated as well as whether individual stimuli can be recalled based on the final state of the network. Since there are 7! = 5040 possible input sequences, we used two different methods to sample this space. In the first we presented a canonical 7-word sequence as well as the 6 sequences obtained by swapping the positions of nearest neighbors. We expected that sequences with nearest-neighbor swaps would be the most difficult to discriminate, so this data set would provide the most stringent measure of sequence discrimination. These 7 sequences serve as a proxy for the full set of sequences obtained by allowing all permutations, but in particular they allow measurement of primacy and recency. In the second method, we generate a list of 70 distinct 7-word sequences obtained using the Latin-squares method such that each stimulus type is presented the same number of times in each serial position across the whole set of sequences. The second method produced more examples of words in a given position so, we expected, would provide a better dataset for position-specific measures such as primacy and recency.

#### 9) Constituent stimulus recognition and probability of first recall

To test which word in a sequence is most likely to be recalled first, we compared the state of the network after presentation of a complete sequence of stimuli to its state after presentation of an individual stimulus on its own. In this test we altered the length of lists, varying the number of stimuli from *N*_*stims*_ = 2 to *N*_*stims*_ = 10, in order to assess whether our networks produced the observed shift from primacy in short lists to recency in long lists.

In a secondary test, we wished to assess the overall likelihood of recall of words in a list. We reasoned that for a word to be recalled, information about its presentation must be present in the final state of the network. As a simple test, we assumed a large vocabulary of 35 words and required that for a word to be recalled, at least in principle, the final pattern of activity in the network should be closer to that word’s individual activity pattern than to any word that had not been presented. Therefore, we measured the activity resulting from each of 35 individual stimuli comprising the vocabulary. We then compare these vocabulary patterns with the final activity after presentation of one sequence of *N*_*stims*_ stimuli. We take the 7 vocabulary patterns which most closely resemble the sequence and consider those corresponding to items which are indeed members of the sequence as the items recalled.

## Results

### Left-Right evidence accumulation and discrimination

Our first goal was to assess whether the network could, following a sequence of six left/right stimuli, produce neural activity capable of distinguishing sequences with more left-stimuli from those with more right-stimuli. Our assessment comprised two tests. In the first test, we produced a confusion matrix, which indicates how distinct are the final activity states following each of the 64 possible combinations of six left/right stimuli. If all 64 stimulus patterns could be encoded distinctly in the network, we hypothesized that appropriate responses to those final activity states could be trained. As seen in Figure 2), perfect distinguishability between final states is possible for some networks. Moreover, for a broad range of parameters, the networks achieve good performance in this test (Figure 3a).

**Figure 3.**
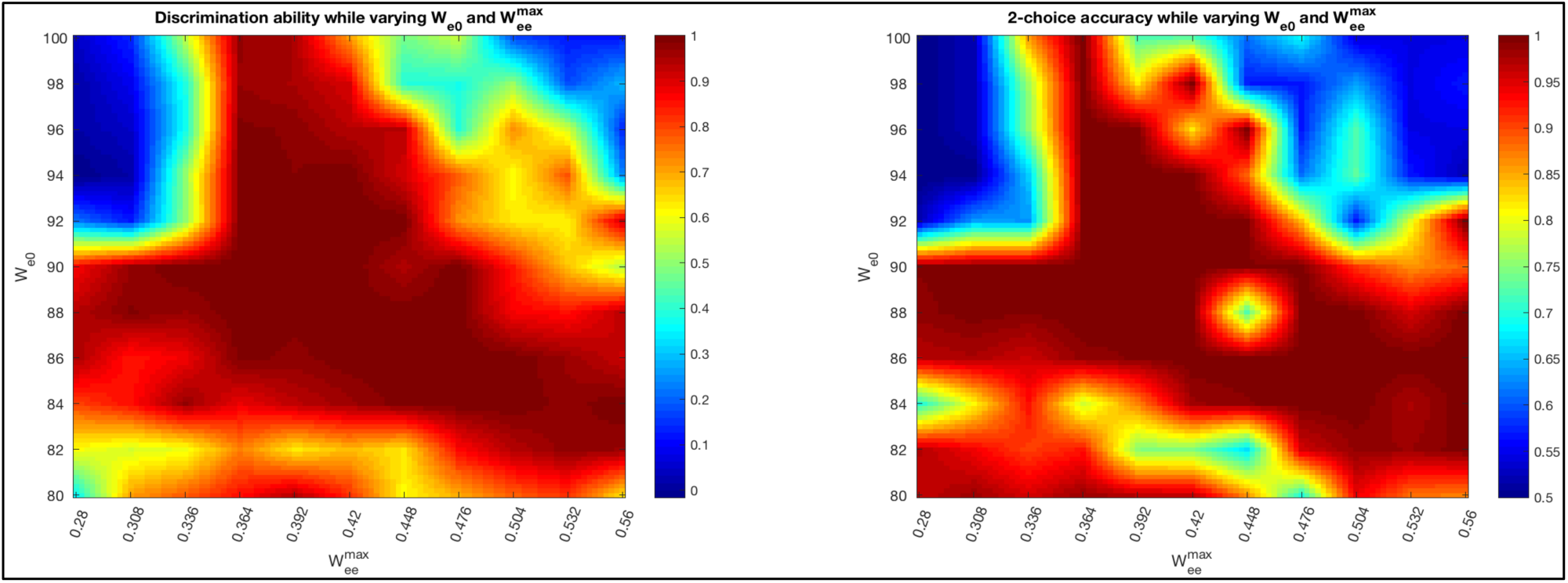
left: Discrimination ability ***κ*** for 121 networks interpolated across the self/cross-excitation space. right: Same as left but with 2-choice decision accuracy plotted for each network (as measured by a perceptron).

For the second test, we trained an output (using a perceptron, see Materials and Methods) to produce a left or right response according to the greater number of left or right stimuli. In this manner, we could compare the response generated by the network with animal behavior. The results in Fig. 3 (right) indicate again that good performance is achieved over a broad range of parameters.

It is revealing to visualize the trajectory of neural activity for two similar sequences by projecting to the space of the first three principal components. That is, we chose as a basis the three axes along which arose the greatest variation of neural activity across the sequences. In Figure 4 (and see supplementary movies 1 and 2) we see that the two trajectories separate and move to distinct attractor states when the stimuli differ at a single point in the sequence. Thereafter, with identical following stimuli, the trajectories remain almost parallel, with a slow decline in the separation between activity states as further identical stimuli are added.

**Figure 4.**
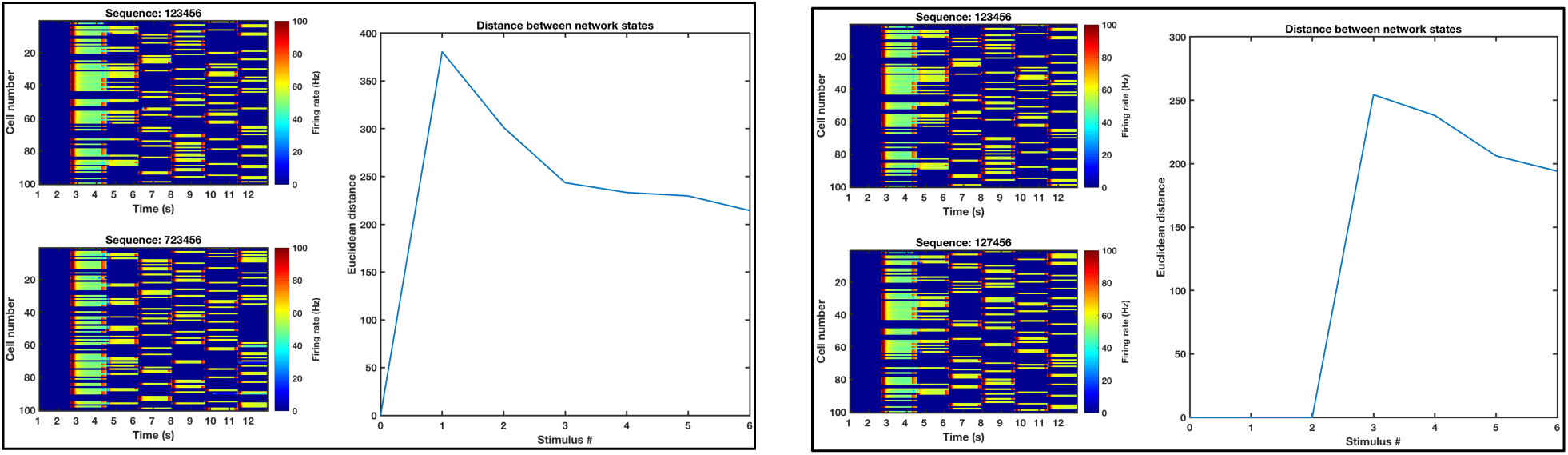
Comparisons of trajectories through activity space due to sequences that differ in only 1 position. a) Networks differ in their 1^st^ stimulus. Left: Network activity versus time. Each row indicates the firing rate of one unit. While successive stimuli cause changes in the activity patterns, those changes are in part determined by the prior activity pattern. Right: Euclidean distance between the network states induced by the two sequences. Since the 2 sequences differ in their first stimulus, the trajectories immediately separate. b) Same as a) except the two sequences compared here differ in their 3^rd^ stimulus.

The successful networks could be characterized as having low noise. A particular question when assessing whether such networks could be operating in the brains of animals is whether they are robust to the levels of firing rate variability typically observed in vivo. Moreover, as performance degrades in these networks – as it must with increased noise – we wished to assess whether the patterns of errors would allow us to make predictions about errors made by animals in behavioral tasks. Therefore, we tested the networks with increased noise amplitude, *σ*.

The most important metrics of network performance for the L/R test are total discrimination ability and evidence accumulation ability. As a secondary feature, we were interested as to whether or not primacy and recency are present, perhaps simultaneously in one network, as such patterns would allow us to make predictions regarding animal behavior. We find that for noise amplitude *σ* = .001 (corresponding to ~1-2Hz noise oscillations) the network is able to discriminate with between 90% – 96% accuracy for a wide range of parameters. In Fig. 5 we plot three two-dimensional slices of both the four-dimensional space of network parameters, as well as the two-dimensional space spanned by the fraction of cells receiving input, and the amplitude of current making up the stimuli. These slices all intersect at a fiducial parameter value

**Figure 5.**
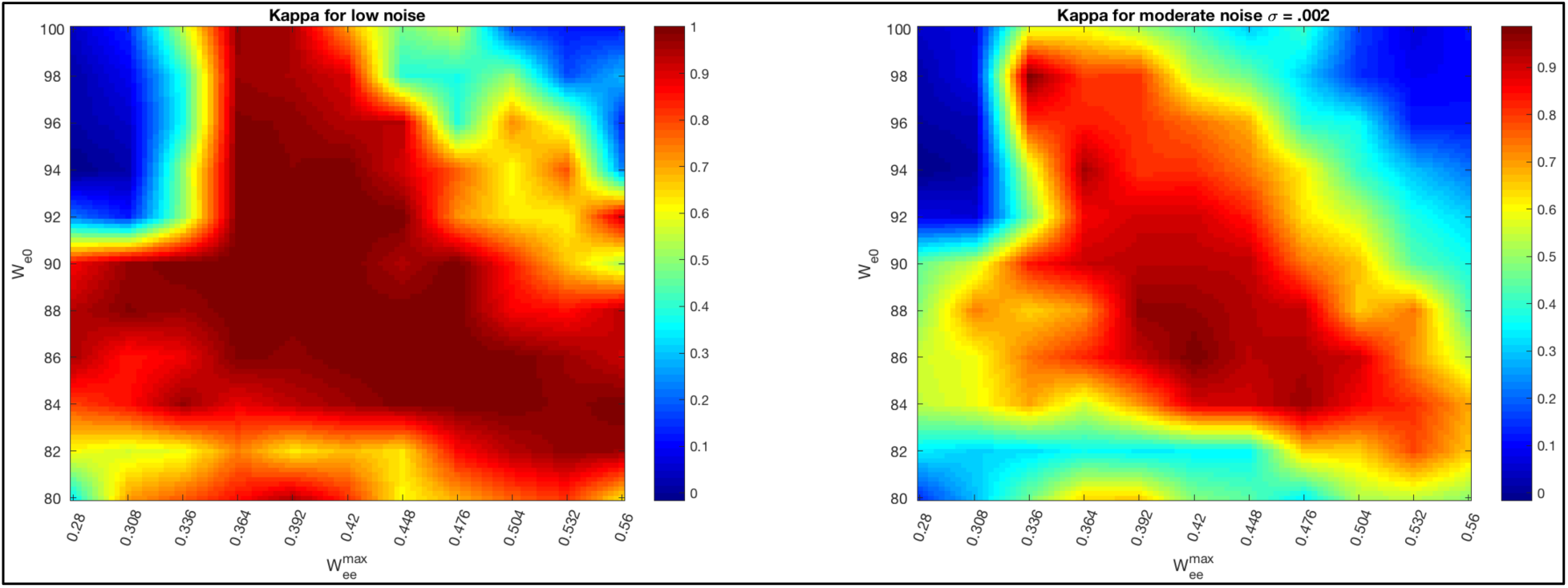
Sequence discrimination ability for networks with very low (**left: σ < .0001**) or moderate noise (**right: σ = .002**).

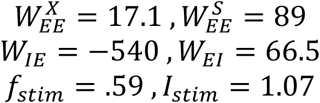

#### Comparison with Morcos & Harvey

We can compare the percentage of left choice vs. net evidence data with actual mouse behavioral data [33]. In Fig. 6, the psychometric curves are taken from those networks in Fig. 3 whose 2-choice accuracy (as measured by a perceptron) was greater than 75%. Note that a perceptron with perfect choice accuracy would produce a step-function as its psychometric curve. We find close agreement in the choice behavior of our network and the 5 mice imaged by Morcos & Harvey. As found in the real mice, network activity did not seem to directly represent different accumulated evidence values, but rather distinct network states dependent on the cue history. In our simulations, the network was started from the same initial conditions (quiescent) each trial. As such, for low noise values, identical sequences led to very similar final network states (as can be seen in the confusion matrix). In the real mice however, it was found that identical stimulus sequences could lead to a diversity of network states. This is likely due to the fact that the mouse’s starting internal state will never be exactly the same from trial to trial and that the network in the mouse’s posterior parietal cortex is likely undergoing plasticity while learning the task.

**Figure 6.**
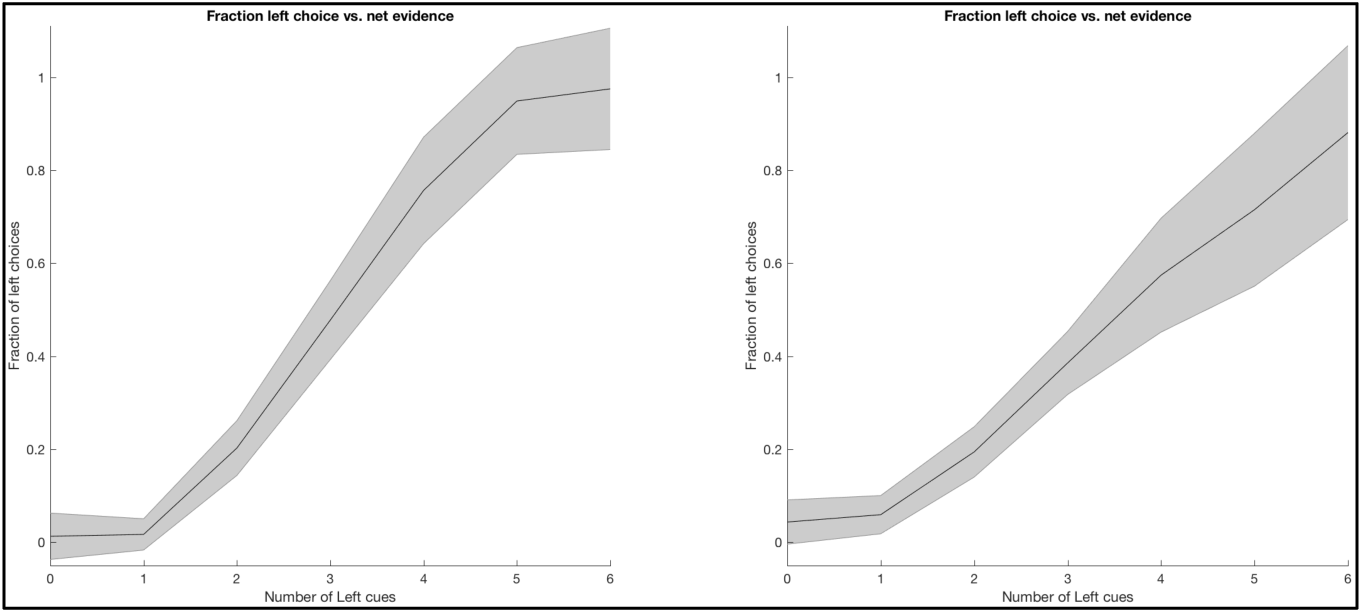
Likelihood of left choice vs. net evidence. Left: Choice curves for very low levels of noise from networks with 2-choice accuracy > .75. Right: Same as left but for higher noise values (σ = .002 corresponding to ∼2Hz fluctuations in firing rate). Shaded region shows standard deviation across networks.

#### Non-repeating sequences of 7 stimuli

The second task we performed is analogous to word recall in humans. We presented 70 different sequences (each with 7 distinct stimuli) to the network and again asked how well the network could discriminate between each sequence as well as whether the location and identity of a given stimulus could be identified solely by the final network state (recall). We found again that a wide range of networks could perfectly discriminate between all 70 sequences (Fig. 7).

**Figure 7.**
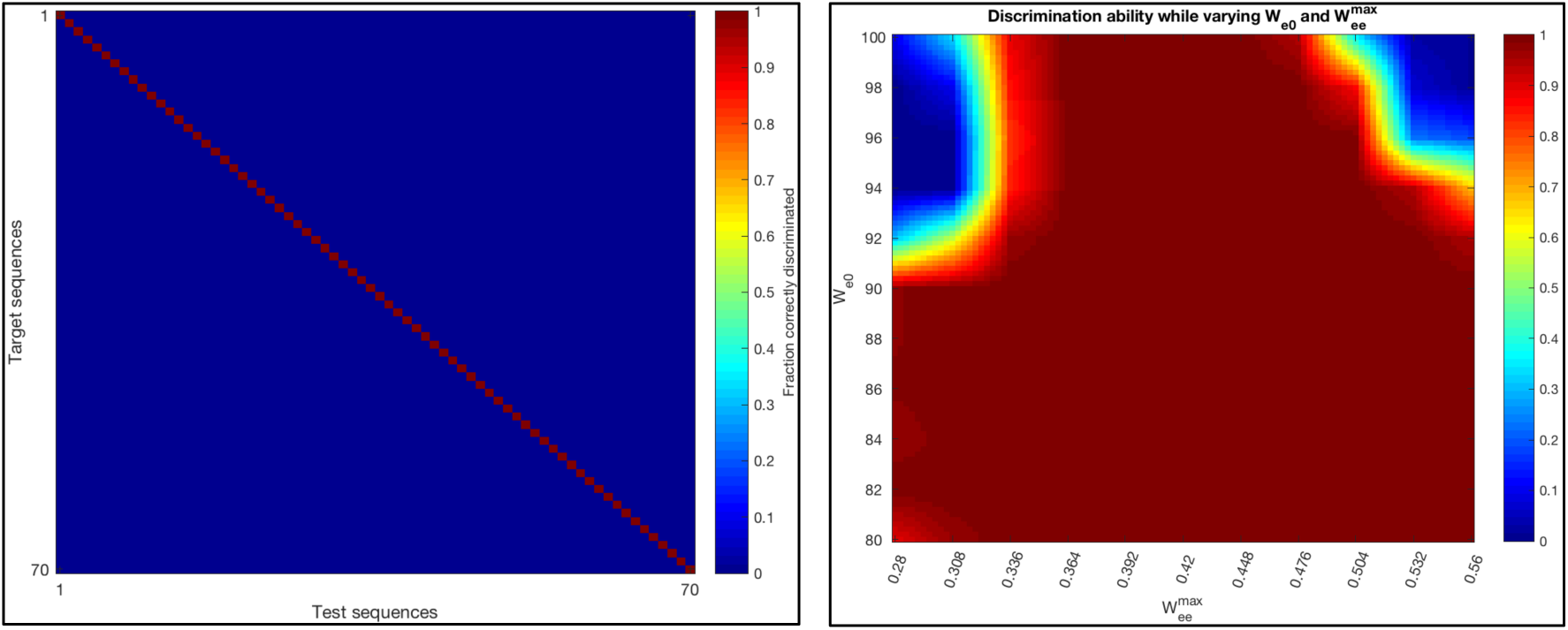
**Left:** Confusion matrix for a network with perfect discrimination in the multi-item task (7 words for 70 distinct sequences). **Right:** Discrimination ability in the 7-word multi-item task for a range of networks in the self/cross-excitation space.

#### Primacy and Recency

In the 7-word task, the network state at the end of the stimulus sequence determines which of the stimuli is first recalled. Therefore, we compared the final network activity pattern to the patterns produced by each of the stimuli alone and took the most similar individual-stimulus pattern to correspond to the stimulus first recalled. We found a significant impacts of stimulus strength and strength of internal network connections. If the stimuli are relatively stronger and the network’s internal connections are relatively weaker, then the network’s activity pattern follows that of later stimuli (more recency) whereas in the opposite condition, more primacy is observed. For a very narrow range of intermediate stimulus strengths networks exhibited both primacy and recency in first recall. When we allowed stimulus strength to vary randomly about a mean along a sequence, the intermediate range of both primacy and recency broadened. Moreover, ubiquitously, as we compared sequences of different lengths we observed a shift from primacy for short lists to recency for longer lists (Figure 8).

**Figure 8.**
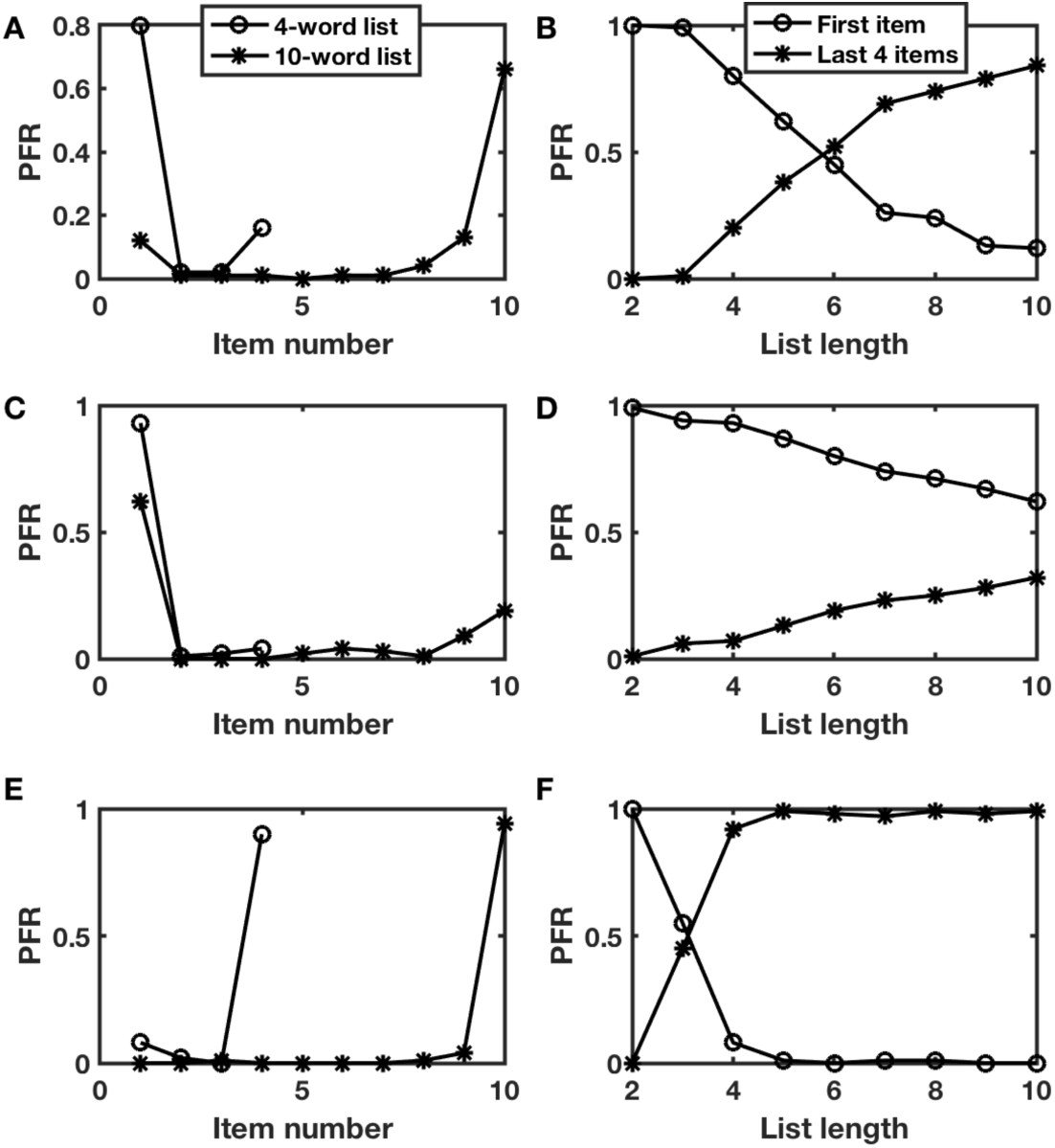
Probability of first recall exhibits primacy in short lists and recency in long lists. **A, B.** With appropriate parameters (***I*_*stim*_** = 0.9 ± 0.2) matches behavioral data. ***C, D.*** Reduced stimulus strength (*I*_*stim*_ = 0.85 ± 0.2) primacy dominates, but still decreases with increased list length. ***E, F.*** Increased stimulus strength (*I*_*stim*_ = 1.0 ± 0.2) recency dominates in all lists beyond length 3. ***A, C, E.*** Probability of first recall (PFR) as a function of item position in the list for 4-item lists (circles) or 10-item lists (asterisks). ***B, D, F.*** Probability of recall for the first item (circles) or up to the last 4 items excluding the first item (asterisks) as a function of list length.

We find that in both tasks, primacy and recency effects are positively correlated (Figures 9-10). For the 2-choice task, networks show a range of primacy/recency scores. The networks that discriminated the best had little or no primacy or recency or actually had negative primacy/recency scores (indicating the opposite of each effect). Some networks were found to show limited recency and inverse primacy (negative primacy scores) however, no networks were found to have the reverse (positive primacy and negative recency, see Figure 9). Additionally, whether considering all networks or just those that discriminate well, both primacy and recency were negatively correlated with discrimination ability (Figure 10).

**Figure 9.**
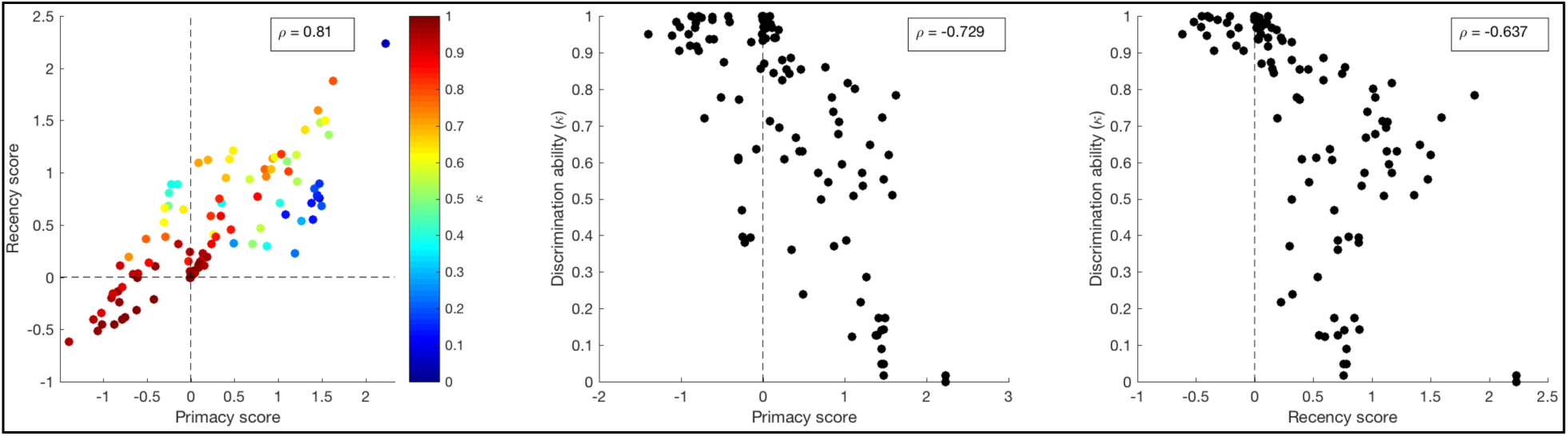
Relationships between primacy, recency, and discrimination ability for the 2-choice task. Data are from the same networks shown in Fig. 3. **Left:** Scatterplot of primacy vs. recency scores. Color indicates discrimination ability **(*κ*). Middle:** Scatterplot of primacy scores vs. discrimination ability. **Right:** Scatterplot of recency scores vs. discrimination ability. Insets: correlation coefficients between each pair of variables.

**Figure 10.**
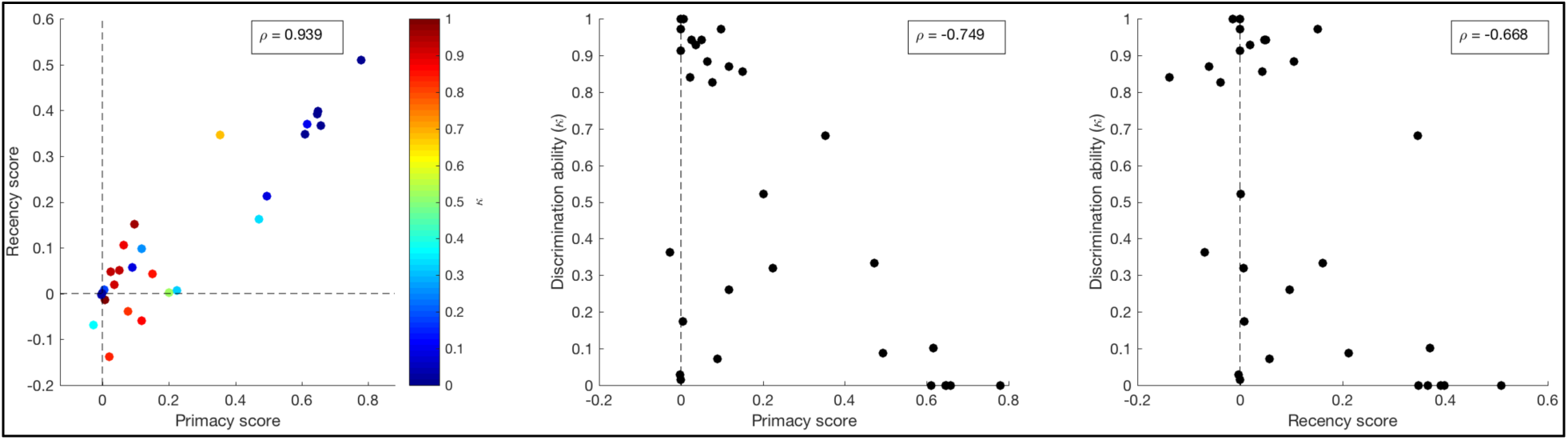
Same as Figure 9, but for the multi-item task (data from networks shown in the right of Fig. 7). Note that the majority of the data points in all 3 plots are overlapping and indistinguishable. In the left plot these overlapping points are at (0,0) while in the middle/right plots they are at (0,1).

Networks with the highest discrimination abilities exhibit neither primacy nor recency (Figure 10). However, networks in a large fraction of parameter space surrounding the optimal value do exhibit both primacy and recency, while still maintaining discrimination ability above 60%. A small fraction of parameter space exhibits both primacy and recency with ability above 90%. It is possible to find networks exhibiting significant primacy and recency, but their discrimination ability is significantly lower. Furthermore, unlike in the 2-choice task, very few of the networks had significantly negative primacy/recency scores.

## Discussion

Our brains allow us to learn to respond appropriately to sequences of events rather than to be merely reactive to each ongoing stimulus. Therefore, circuits of neurons which can produce activity patterns containing information about the temporal pattern of prior stimuli have been an active area of research in computational neuroscience. Particularly powerful have been networks derived from liquid state machines [38, 39], in which a stimulus produces a temporary, decaying trace of neural activity, but which trace interacts with later stimuli and can be extracted by training an appropriate “readout”. More recent work has shown how connections within such circuits can be trained to enable the network to learn almost any stimulus-response mapping [40–42], even in a manner that is robust to noise and time-warping [43, 44].

Our work also follows the paradigm of the neocortex containing neural circuits that can serve as general-purpose computational machines. When the neural activity produces high-dimensional discrete attractor states, which are inherently robust and stable, an appropriate output can be readily trained through reinforcement learning, especially when just a binary response is necessary [3, 45–48]. Therefore, while point-attractor networks lack the dynamic versatility of those based on chaotic systems, so would be less suitable as a basis for motor output, they may be more easily trained for complex cognitive tasks [49], which inevitably require short-term memory [50].

### Pattern separation versus pattern completion

One can think of the two experimental tasks considered in this work as testing the two distinct, desirable, but naturally competing properties of attractor networks—namely, pattern separation and pattern completion [51]. In the T-maze task, pattern separation is required, as distinct responses must be made between patterns of activity generated by an excess of left-stimuli and patterns of activity generated by an excess of right-stimuli. If each possible stimulus sequence produces a distinct final pattern of neural activity, and if the neural activity has high enough dimensionality, then distinct responses to any subset of stimulus patterns can be trained. Performance improves if the network maps differences in stimuli to increasingly different patterns of neural activity.

However, in a recall task, the final activity of the network must regenerate the stimulus sequence that led to it. In the simplest scenario that we test here, we require that the final network state has greater overlap than chance with the state produced by a stimulus alone for that stimulus to be recalled. Such overlap is optimized if successive stimuli do not cause the network’s activity to change by much from its prior state. Therefore, it is perhaps unsurprising that we observed a negative correlation across successful networks, with those performing best in the sequence discrimination task fairing less well in the recall task and vice versa.

For example, by reducing the input fraction *f*_*stim*_, the inhibition never grows enough to shut off any active units and recognition ability approaches unity at the cost of discrimination ability. Similarly, the optimal network for discrimination fares poorly for recognition (Supporting Information S1 Figure). However, this picture changes when we move away from the optimal region of parameter space, to networks scoring below 90% in both categories. The abilities can be positively correlated, since the requirement for multiple, reachable attractor states is common to the two tasks and is not met under many parameter combinations (Supporting Information S2 Figure).

### First recall and overall recall

The first item recalled following a list of stimuli must be generated by the activity in the network that remains after presentation of all stimuli. While one may think that the most recent item would always have the strongest effect on the final activity state, there are broad regions of parameter space where the first item, which imprints on a less active network, causes certain activity patterns that are hard to shift. As the sequence of stimuli grows, more and more changes in the initial privileged state accrue until the final network state becomes more like later than like the first state. Such behavior is observed with changing list lengths in human free recall data. However, in our simulations, in those cases in which long sequences left the final network state closest to that induced by the first stimulus, the network retained little information about late stimuli and was indeed in some cases not influenced by them. Thus, at face value, in 10-word lists when the first item recalled is the first word, then only a few of the other early items would be recalled and none of the later ones. Such behavior would be intriguing if extracted from the human behavioral data, but we suspect it is unlikely because of the absence of many important features in this initial demonstration of the behavior of a randomly connected circuit.

For example, in this paper we do not include correlational synaptic strengthening. Such strengthening is undoubtedly important in linking together the representations of successively presented words to form the observed associations which would aid their recall [52–54]. Such information, which lies outside of the final state of activity in the network as used in our assessments here, is most likely essential for list recall in the multiple-item presentation task.

### Impact of noise

Even in the absence of noise, the networks’ ability to discriminate between sequences of six or seven stimuli with minimal variations in the sequences is non-trivial. While distinct stimuli cause trajectories to diverge in chaotic networks, the history-dependence of the trajectory of activity in attractor networks has received less widespread attention. Whereas chaotic networks are naturally extremely sensitive to noise fluctuations, so careful attention to their robustness to such fluctuations can be necessary when training them to solve tasks, point attractor states are inherently robust to noise fluctuations up to the size of the barrier between attractor states. In our network the size of the barrier corresponds to a fluctuation in firing rate sufficient to render an active unit quiescent or a quiescent unit active. Therefore, we investigated the robustness of sequence discrimination to noise fluctuations in our networks.

In the point-attractor networks, the activity states remain relatively stable between stimuli and noise has its strongest impact during a stimulus when the system switches from one state to one of the many other states nearby in high-dimensional space. At such switching times, the network is much more sensitive to noise fluctuations as the barriers separating the zero-noise trajectory from trajectories to nearby attractor states is much smaller than the barriers between the attractor states themselves. Yet, even without training (and alterations of connection strengths), we did find that some networks were capable of producing good performance—at least at the human level—even while exhibiting considerable variability in their firing rates (e. g. 10Hz fluctuations during the delay periods).

## Acknowledgements

We are grateful to the Swartz Foundation for supporting this research (Grant 2016-6) and for funding from the NIH under grants R01DC009945 (from NIDCD) and R01NS104818 (from NINDS). This work is the responsibility of the authors and is not necessarily endorsed by NIH.

